# Epitope spreading of Lyme autoantigen apoB-100 and CD4+ T cell responses to *Borrelia burgdorferi* Mcp4 are regulated by IL-10 in murine Lyme disease

**DOI:** 10.1101/2023.06.16.545225

**Authors:** Rebecca Danner, Michaela Pereckas, Joseph R Rouse, Amanda Wahhab, Lauren Prochniak, Robert B Lochhead

## Abstract

*Borrelia burgdorferi*, the causative agent of Lyme disease (LD), has evolved immune evasion mechanisms to establish a persistent infection in their vertebrate hosts, resulting in chronic inflammation and autoimmune T and B cell reactivity in many *B. burgdorferi*-infected individuals. In this study, we used an unbiased immunopeptidomics approach to identify foreign and self MHC class II peptides isolated from inguinal and popliteal lymph nodes from *B. burgdorferi*- infected C57BL/6 (B6) mice, which develop mild, self-limiting LD; and from infected B6 Il10^-/-^ mice, which develop severe, persistent LD. Nearly 10,000 MHC-II peptides were identified by LC-tandem MS analysis which included many peptides derived from proteins abundant in arthritic joints that are associated with inflammation, tissue repair, and extracellular matrix remodeling. Notably, the number and variety of unique peptides derived from apolipoprotein B- 100 (apoB-100); a validated autoantigen in human Lyme arthritis (LA), atherosclerosis, and liver disease; was greatly expanded in lymph nodes of infected mice, particularly in Il10^-/-^ mice at 4 weeks (6-fold increase) and 16 weeks (15-fold increase) post-infection, compared with uninfected mice, indicating epitope spreading. One of the apoB-100 peptides identified in infected, but not uninfected, B6 and Il10^-/-^ mice was APOB_3500-3515_, an immunogenic cryptic epitope in murine autoimmune atherosclerosis. No apoB-100 peptides had sequence homology to any *B. burgdorferi* antigens. Surprisingly, only six peptides derived from *B. burgdorferi* proteins were validated in this study. One of these *B. burgdorferi* epitopes, derived from methyl-accepting chemotaxis protein Mcp4 (BB0680), was an immunogenic target of CD4+ T cell responses in *B. burgdorferi*-infected Il10^-/-^ mice, but not in B6 mice. In conclusion, this study has shed light on the importance of IL-10 in suppressing epitope spreading and limiting *B. burgdorferi*-specific CD4+ T cell responses. Furthermore, this study supports epitope spreading and exposure of cryptic antigens as likely mechanisms of infection-induced apoB-100 autoimmunity in LD.

**AUTHOR SUMMARY:** Lyme disease is caused by infection with the spirochetal pathogen Borrelia burgdorferi, and affects ∼500,000 individuals in the U.S. annually. T cell responses to both host and pathogen are dysregulated during infection, resulting in chronic infection and frequent development of autoimmunity. To assess the immune-relevant CD4+ T cell epitopes presented during development of Lyme disease, we used an unbiased, immunopeptidomics approach to characterized the MHC class II immunopeptidome in mice infected with *Borrelia burgdorferi*. We identified nearly 10,000 unique peptides. Peptides derived from apoB-100, a known human Lyme autoantigen, were highly enriched in infected mice, compared with uninfected controls, and showed evidence of epitope spreading. Furthermore, we identified several peptides derived from *Borrelia burgdorferi*, including one immunogenic peptide from a methyl-accepting chemotaxis protein, Mcp4. Interestingly, both apoB-100 epitope spreading and immune responses to Mcp4 were observed in mice lacking the anti-inflammatory cytokine IL-10, indicating an important role of IL-10 in suppressing T cell responses to Mcp4 and epitope spreading of Lyme autoantigen apoB-100.

## INTRODUCTION

Autoimmune disorfers impact ∼7% of the U.S. population and are increasing in frequency(1, 2). Loss of immune tolerance during a microbial infection is thought to be a common autoimmune trigger for many chronic inflammatory and autoimmune diseases(3). However, identifying a causal relationship between an infection and subsequent autoimmunity remains an enormous challenge. Lyme disease (LD) is an example of infection-induced autoimmunity where the autoimmune trigger — infection with the tick-borne spirochete, *B. burgdorferi* — is known with certainty(4). The purpose of this study was to identify foreign and self MHC class II (MHCII) peptides presented by antigen presenting cells (APCs) of mice infected with *B. burgdorferi* to elucidate the mechanisms of T cell responses to *B. burgdorferi* and development of LD-associated autoimmunity.

LD is a global publich health concern(5) and is the most common vector-borne disease in North America, with ∼500,000 cases per year(6). In the U.S., LD is transmitted by infected adult- or nymph-stage Blacklegged ticks, or *Ixodes scapularis*(7), and is an infection-induced, multi- system disorder that affects the skin, heart, joints, or neurologic tissue(8, 9). Early LD symptoms include erythema migrans rash, fever, chills, headache, fatigue, myalgia, arthralgia, and swollen lymph nodes(8, 10). Late-stage symptoms include severe joint pain and swelling, carditis, facial palsy, meningitis, nerve pain, numbness, and dizziness(8). Lyme arthritis (LA), the most common late-stage disease manifestation, is usually effectively treated with antibiotic therapy(11). In ∼10-20% of cases, however, their arthritis can persist or worsen for months or years following antibiotic therapy and apparent spirochetal killing(11); a condition known as post-infectious, or post-antibiotics, LA(4). In both post-infectious LA and in a subset of patients with other late-stage disease manifestations, called posttreatment Lyme disease syndrome (PTLDS)(12, 13), additional courses of antibiotics are not effective at resolving persistent symptoms(14, 15). In the case of post-infectious LA, use of disease-modifying antirheumatic drugs are typically effective at resolving joint inflammation, similar to treatments used for rheumatoid arthritis (RA), the prototypical autoimmune joint disease(11). As in RA, the post-infectious LA synovial lesion is characterized by marked synovial hyperplasia and is frequently accompanied by autoimmune T and B cell responses against self-antigens derived from extracellular matrix proteins and proteins involved in vascular damage and repair(4, 16–19). However, due to a paucity of animal models, the mechanisms of infection-induced autoimmunity in LA are incompletely understood.

Despite a robust antibody response, *B. burgdorferi* persist within their vertebrate hosts for months or years through manipulation of host innate and adaptive immune responses(20–22). LD spirochetes are capable of subverting adaptive T cell responses during infection(20, 23). This T cell suppression is likely mediated through multiple mechanisms such as down-regulation of MHC class II molecules(24), upregulation of the inhibitory PD-1/PD-L1 pathway(25), and altered effector function during infection(26). However, no CD4+ T cell antigens specific for *B. burgdorferi* proteins have been identified in mice infected with *B. burgdorferi*, which has limited our ability to fully understand these immune evasion mechanisms(25).

## RESULTS

### Analysis of the murine Lyme arthritis MHC II immunopeptidome

For this study we used two established mouse models of Lyme disease that recapitulate the range of LA severity and duration seen in humans(4). When infected with *B. burgdorferi*, C57BL/6 (B6) mice develop mildly inflammatory LA that peaks at approximately 4 weeks post-infection, then resolves over several weeks, although the infection itself persists indefinitely(27). In contrast, Il10^-/-^ mice infected with *B. burgdorferi* develop severe, persistent LA that is driven by dysregulated Th1 responses that persist for at least 16 weeks, despite these mice harboring ∼10-fold fewer bacteria compared with infected B6 mice(28–30), similar to autoimmune- promoting conditions seen in patients with post-infectious LA(4).

To identify the repertoire of MHC class II peptides presented by APCs during infection, we infected B6 and Il10^-/-^ mice with 2 x 10^3^ *B. burgdorferi* (strain N40, a generous gift of Dr. Mark Wooten) or BSKII media alone for 4 or 16 weeks (22-23 mice per group, **Fig 1A**). Inguinal and popliteal lymph nodes were harvested and pooled, peptide-MHC-II complexes were isolated by immunoaffinity purification, and peptides were eluted from the MHC-II molecules with formic acid, as previously described(31). Eluates were analyzed by LC-MS/MS using a Thermo Orbitrap Fusion Lumos (see Supplementary tables for analysis settings). Mass spectrometry data was processed using ProteomeDiscoverer 2.4 and validated using Percolator and Protein FDR validator to identify candidate peptides derived from mouse or *B. burgdorferi* (see Supplementary material for data processing pipeline). To account for possible errors in genome annotation, both the B31 type strain and N40 strain proteomes were analyzed.

**Figure 1:**
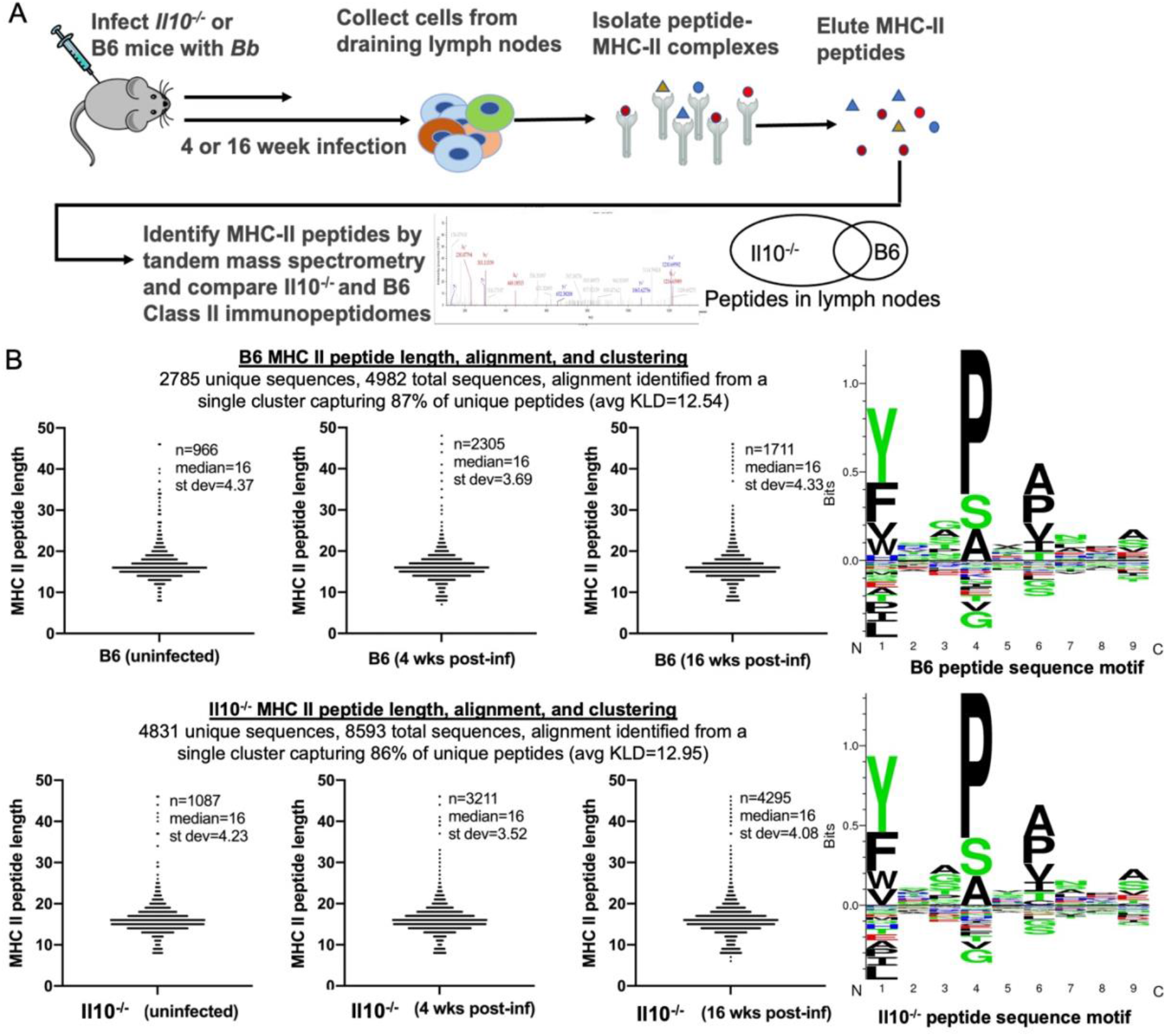
Analysis of MHC-II peptides isolated from lymph nodes of Bb-infected B6 and Il0^-/-^ mice. (A) Experimental design of the current study is shown. (B) MHC class II peptide length, alignment, and clustering analysis of epitopes identified by LC/MS/MS from pooled inguinal and popliteal lymph nodes from C57BL/6 (B6) and B6 Il10^-/-^ mice infected with *Bb* for 0, 4, or 16 weeks (22-23 mice per group). The number, median length, and standard deviation of peptide length are shown for each group of mice. Peptide sequence motifs were generated using Seq2Logo 2.0 (TDU Health Tech), and the cluster capturing the largest percentage of unique peptides is shown for each mouse strain.

We identified 4,982 candidate peptides (2,785 unique sequences) from B6 mice and 8,593 candidate peptides (4,831 unique sequences) from Il10^-/-^ mice (**Fig 1B, Supplementary File**). The MHC II peptide length, alignment, and sequence binding motif clustering were similar between mouse strains for all groups, and the median peptide length was 16 amino acids (**Fig 1B**), the typical size of MHC II bound peptides.

In uninfected B6 and Il10^-/-^ mice, we identified ∼1,000 peptides in each group, predominantly consisting of proteins involved in antigen uptake, processing, and presentation (**Supplementary File**). However, we observed marked differences between mouse strains at 4 weeks and 16 weeks post-infection. There were more peptides identified in 4 week- versus 16 week-infected B6 mice, and more peptides identified in 16 week- versus 4 week-infected Il10^-/-^ mice, consistent with mild and self-limiting LA in B6 mice, versus persistent and T cell-mediated LA in Il10^-/-^ mice.

KEGG analysis revealed a number of similarities and differences between strains in the array of proteins presented in lymph nodes, likely reflecting changes in the protein microenvironments in both the lymph nodes and in nearby joints (**Table 1**). All infected groups contained a large number of peptides derived from proteins involved in metabolism and tissue damage and remodeling, although there was marked variability between strains at the two time points. All infected groups had enrichment of proteins involved in metabolism of amino acids, pyruvate, carbon, and/or cholesterol; whereas only infected Il10^-/-^ mice contained a high number of peptides derived from proteins involved in the TCA cycle. Proteins involved in leukocyte trans- endothelial migration, also involved in *B. burgdorferi* migration into the ECM, were over-represented in both B6 and Il10^-/-^ mice infected for 4 weeks, but only in Il10^-/-^ mice at the 16 weeks post-infection time point. Platelet activation proteins were enriched in B6 mice at 4 weeks post-infection, and in Il10^-/-^ mice at 16 weeks post-infection. Proteins involved in ECM-receptor interactions, indicative of tissue repair and remodeling, were enriched in B6 mice infected for 16, but not 4, weeks; but in Il10^-/-^ mice infected for 4, but not 16, weeks.

**Table 1.**
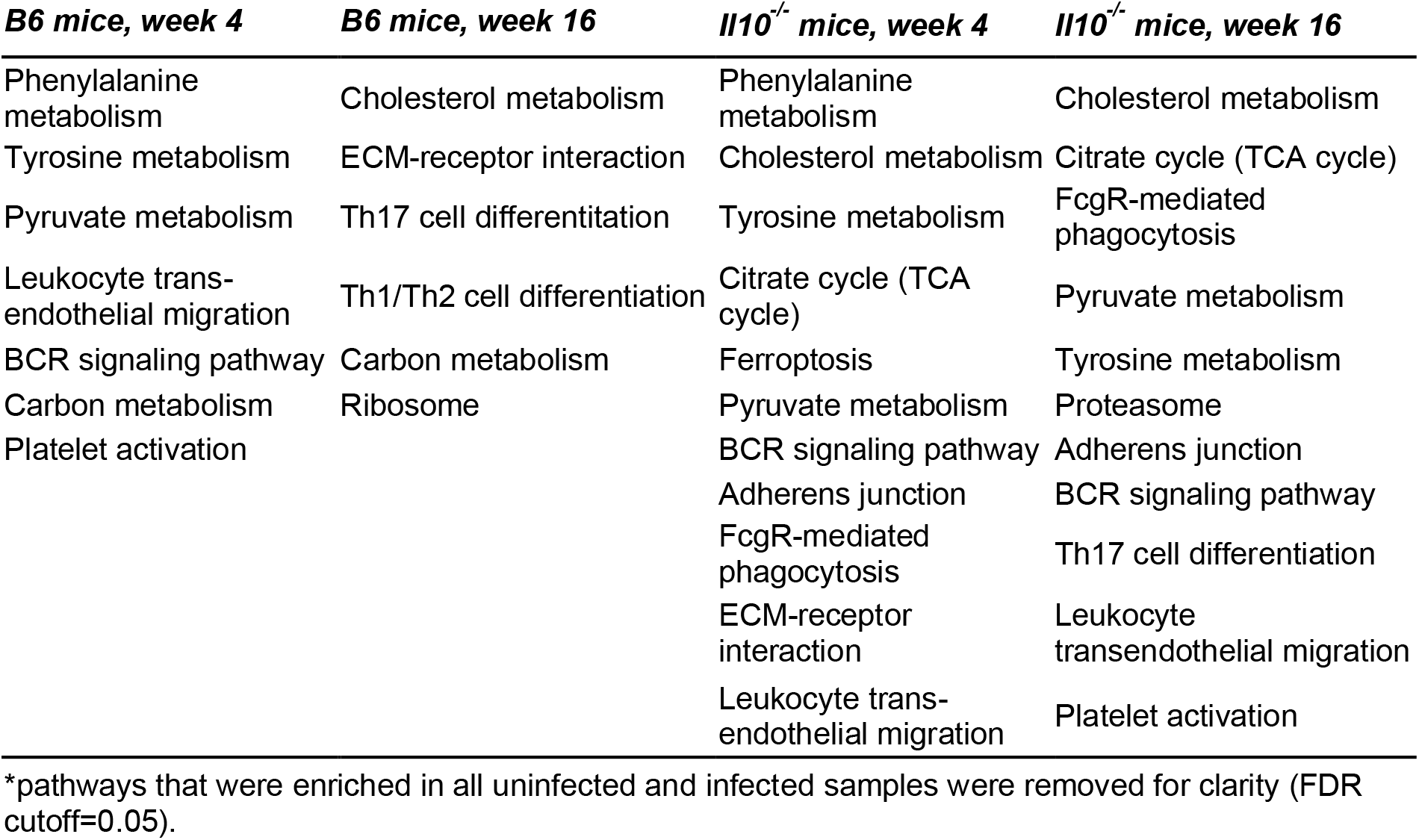
KEGG pathways* enriched in pools of MHC II peptides isolated from *Bb*-infected B6 and Il10^-/-^ mice.

All groups also showed enrichment of proteins involved in B and T cell responses, but there were some notable differences between strains. For example, proteins involved in B cell responses were enriched in B6 mice infected for 4 weeks, but not 16 weeks, and in Il10^-/-^ mice at both time points. However, proteins involved in FcgR-mediated phagocytosis were only enriched in infected Il10^-/-^ mice, but not B6 mice, at both time points. Proteins involved in T cell responses, particularly Th17 responses, were over-represented in both B6 and Il10^-/-^ mice at 16 weeks, but not 4 weeks, post-infection.

### Epitope spreading of apoB-100 peptides during *Borrelia burgdorferi* infection

Enrichment of proteins involved in cholesterol metabolism (**Table 1**) was of particular interest to us for numerous important reasons. First, there is an association between hypercholesterolemia, hyperlipidemia, and liver abnormalities with increaded risk of LD, more severe LD symptoms during early infection, and development of PTLDS following antibiotic therapy(32–35). Second, apolipoprotein B-100 (apoB-100), the dominant lipoprotein of low-density lipoprotein (LDL) particles responsible for cholesterol transport from the liver to peripheral tissues(36), is a T cell autoantigen in ∼50% of patients with LA(16). Interestingly, apoB-100 T cell autoimmunity is also present in vascular inflammation and atherosclerosis(37–49), and following liver damage(50) (see **Table 2** for a summary). Third, LDL particles are strongly linked to vascular inflammation, which is common in numerous tissues in LD, including skin(51), hearts(52, 53), joints(53, 54), and the central nervous system(55). Fourth, *B. burgdorferi* membrane cholesterol, glycolipids, and phospholipids are acquired from their mammalian hosts(56–59), and are potent immunogens in LD(58, 60–62).

**Table 2:** MHC class II epitopes from apoB-100 and their associated T cell and disease phenotypes*.

| *Peptide ID* | *Associated T cell phenotype* | *Reference no.* |
| --- | --- | --- |
| APOB<sub>655-669</sub> | CD4+ T cell immunogen in human LA, Th1 | Ref. 16 |
| APOB<sub>790-810</sub> | CD4+ T cell immunogen in human LA, Th1 | Ref. 16 |
| APOB<sub>978-992</sub> | CD4+ T cell immunogen in mouse CVD, Treg/Th1/Th17 | Refs. 38, 43, 44 |
| **APOB<sub>1942-1957</sub>** | Unknown | <u>Our study</u> |
| APOB<sub>2026-2044</sub> | CD4+ T cell immunogen in human LA, Th1 | Ref. 16 |
| APOB<sub>2214-2233</sub> | CD4+ T cell immunogen in human LA, Th1 | Ref. 16 |
| **APOB<sub>2919-2934</sub>** | Unknown | <u>Our study</u> |
| APOB<sub>3030-3044</sub> | CD4+ T cell immunogen in mouse and human CVD and NAFLD, Treg/Th1/Th17 | Refs. 45, 46, 49, 51 |
| **APOB<sub>3066-3082</sub> (P1#)** | MHC-II binder, does not induce CD4+ T cell activation in mice | <u>Our study</u>, Ref. 44 |
| APOB<sub>3136-3155</sub> | CD4+ and CD8+ T cell immunogen in mouse CVD, Treg | Refs. 48, 50 |
| **APOB<sub>3500-3515</sub> (P2<sup>#</sup>)** | CD4+ T cell immunogen in mouse CVD, Treg | <u>Our study</u>; Refs. 43, 44 |
| **APOB<sub>3807-3820</sub>** | Unknown | <u>Our study</u> |
| APOB<sub>4411-4430</sub> | CD4+ T cell immunogen in human CVD and NAFLD, Th17 | Ref. 51 |

\* LA= Lyme arthritis, CVD=cardiovascular disease, NAFLD=nonalcoholic fatty liver disease, # See Figures 2 & 3

Because of these numerous links between LDL particles, apoB-100, and T cell autoimmunity in LD, we analyzed MHC II presented peptides derived from apoB-100 during *B. burgdorferi* infection more carefully. Despite being over 4,500 amino acids long, we were only able to detect one unique apoB-100 peptide (P1) in lymph nodes from uninfected B6 and Il10^-/-^ mice (**Fig 2**).

**Figure 2.**
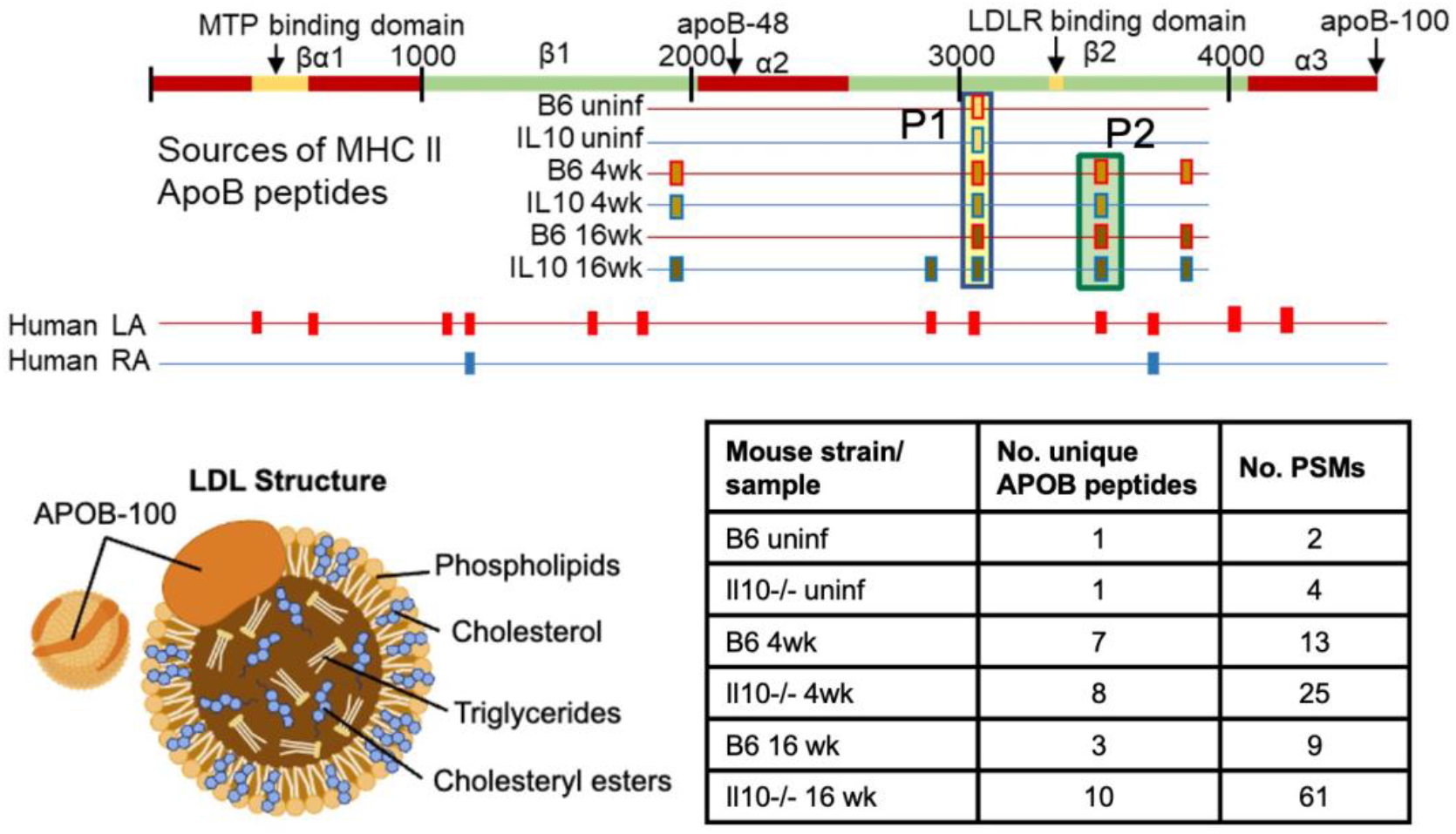
Epitope spreading of apoB-100 MHC II peptides during *Bb* infection. APOB protein is represented in linear form with its five major domains, two isoforms (apoB-48 and apoB-100), microsomal triglyceride transfer protein (MTP), and LDL receptor (LDLR) binding domains labeled. Also shown is a cartoon depiction of the LDL structure. Depicted are approximate locations of MHC II peptides identified in B6 or Il10^-/-^ mice at 0 (uninf.), 4 or 16 weeks post-infection; or in human post-infectious LA or RA synovial tissue mined from previously published data. The epitope highlighted in the yellow box is the non-immunogenic peptide P1 corresponding to APOB_3066-3082_, and the epitope highlighted in the green box is peptide P2 corresponding to APOB_3500-3515_ (see Table 2). The inserted table contains a summary of the number of unique APOB peptides and the number of peptide-spectrum matches (PSMs) for each peptide pool. MHC II peptides from human LA and RA synovial tissue were retrieved from supplemental data from Wang et al (ref. 65).

This P1 peptide has been shown by others to be a strong MHC-II binder in B6 mice, but does not induce CD4+ T cell activation, indicating that P1-reactive T cells are efficiently cleared in the thymus during negative selection(43). However, in mice infected with *B. burgdorferi* for 4 weeks, the number of unique APOB peptides identified increased from 1 to 7 in B6 mice, and from 1 to 8 in Il10^-/-^ mice (**Fig 2**). One of the peptides only detected in infected mice, labeled P2, is known to elicit CD4+ T cell responses in mice, and activated P2-reactive T cells produce high levels of IL-10 and have an anti-inflammatory effect on atherosclerosis(42, 43), indicating that P2 is a cryptic antigen and P2-reactive T cells undergo incomplete negative thymic selection. At 16 weeks post-infection, we observed a contraction of unique APOB peptides from 7 to 3 in B6 mice, but continued expansion of APOB from 8 to 10 unique peptides, and from 25 to 61 peptide-spectrum matches (PSM), in Il10^-/-^ mice (**Fig 2**). These data suggest that *B. burgdorferi* infection induces apoB-100 epitope spreading(63) that is negatively regulated as inflammation resolves in B6 mice in an IL-10 dependent manner, but continues unabated in mice lacking IL- 10.

Evidence of apoB-100 epitope spreading was also observed in a published immunopeptidomics dataset from human LA and rheumatoid arthritis (RA) patients(64). This dataset identified 12 sets of APOB peptides in synovial tissue from human post-infectious LA patients, but only 2 sets of APOB peptides in synovial tissue from RA patients(64). None of the apoB-100 peptides identified in mice or humans showed sequence homology to any *B. burgdorferi* proteins (not shown), which argues against the molecular mimicry hypothesis of infection-induced autoimmunity(65).To our knowledge, apoB-100 autoimmunity has not been reported in patients with RA.

### Localization of apoB-100 peptides within protein structure

Mapping of murine MHC-II presented apoB-100 peptides onto the *in silico* human model of the parent protein revealed that very few of the amino acids found in mice were unmatched to the human sequence (**Figure 3**). The secondary structure of the indentified peptides revealed that all but one contained unstructured regions flanked by beta-sheets. While beta-sheets tend to be hydrophobic, the intervening unstructured loops would be easily cleavable by proteases while being processed within the reducing environment of the phagolysosome, which could further denature the protein and render areas more accessible to cleavage. In addition to this, the inflammatory microenvironment surrounding the site of *B. burgdorferi* infection could change the overall conformation of apoB-100 due to differences in pH and secreted effector proteins.

**Figure 3.**
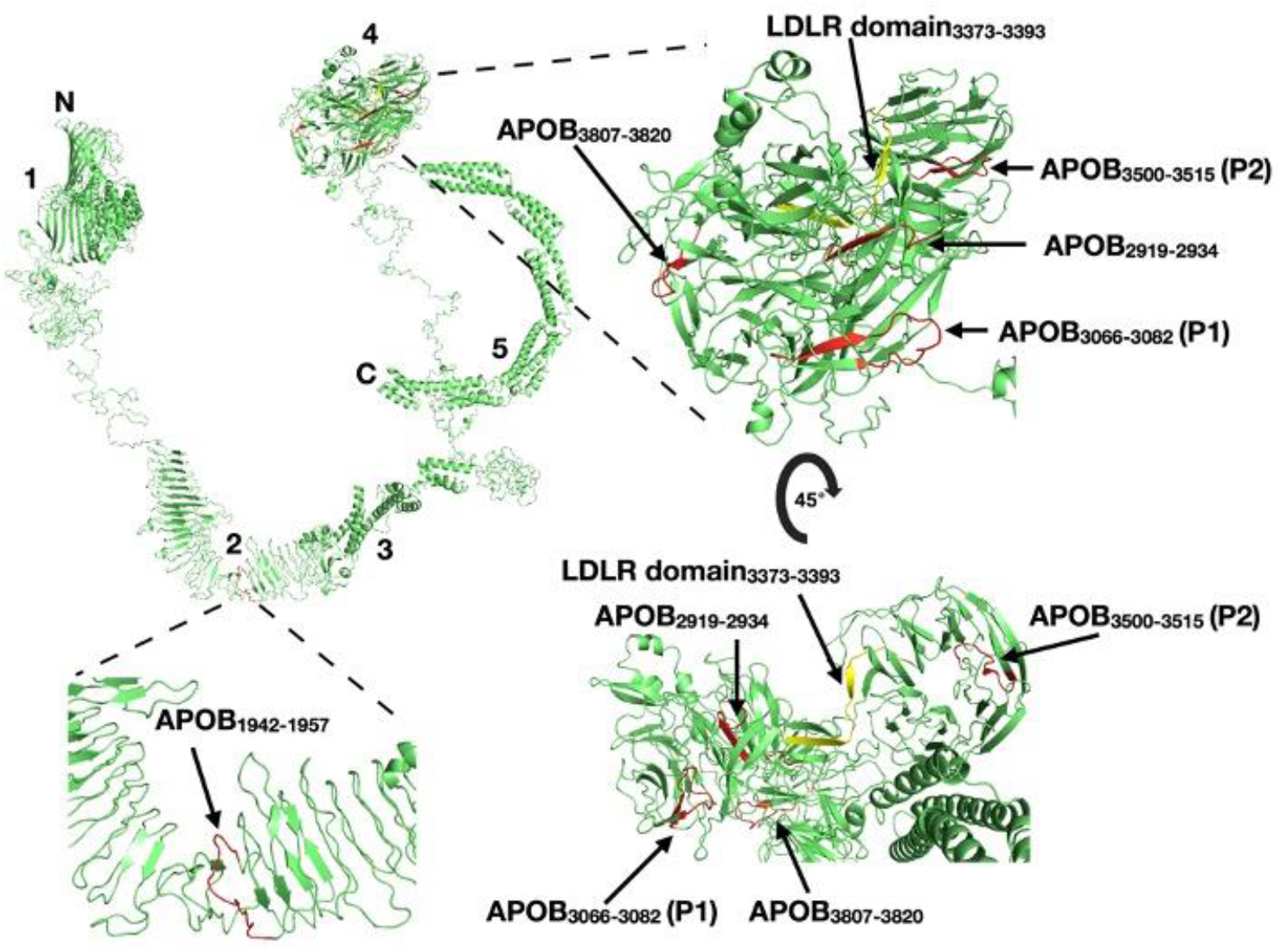
MHC-II presented apoB-100 peptides mapped to whole protein. Murine apoB-100 peptides identified via LC-MS/MS were mapped back to the *in silico* generated model of the total human protein. P1 and P2 indicate peptides shown in Fig. 2. Numbers indicate the five domains of apoB-100 shown in Fig. 2 (1=N-terminal β⍺1 domain containing the MTP binding domain, 2=β1 domain, 3-⍺2 domain, 4=β2 domain containing the LDLR domain, 5=C-terminal ⍺3 domain). Four of the five unique peptides identified (red) occur within the β2 domain of apoB-100 and contain stretches of unstructured loops flanked by beta-sheets. The LDL receptor binding domain (yellow) also lies within the 2 domains.

Moreover, 80% of the peptides identified in this study were located within the same β2 domain as the LDL receptor (LDLR) binding motif. LDLR is expressed by all immune cells, including APCs, which facilitates binding and internalizion of the apoB-100 coated LDL particle(66, 67). Interestingly, hypercholesteremic LDLR KO mice have endogenous autoreactive T cells against the LDLR binding motif of apoB-100(38).

### Synovial inflammation, neovascularization, and fibrosis in infected mice

In human LA, apoB-100 autoimmunity is associated with neovascularization of the synovial sublining and accumulation of activated fibroblasts(16). Infected Il10^-/-^ mice had significantly increased synovial hyperplasia surrounding the tibiotarsal tendon, compared with uninfected Il10^-/-^ mice, and both infected and uninfected B6 mice (**Fig 4A**), consistent with previous studies(28, 52). Interestingly, infected Il10^-/-^ mice, but not B6 mice, have evidence of neovascularization within the inflamed synovium, as measured by CD31 (PECAM) staining (**Fig 4B**). Analysis of collagen deposition and fibrosis using Masson’s trichrome staining revealed evidence of ECM remodeling in the synovial lining of both B6 and Il10^-/-^ mice (**Fig 4C**), which was not significantly different between the two strains.

**Figure 4.**
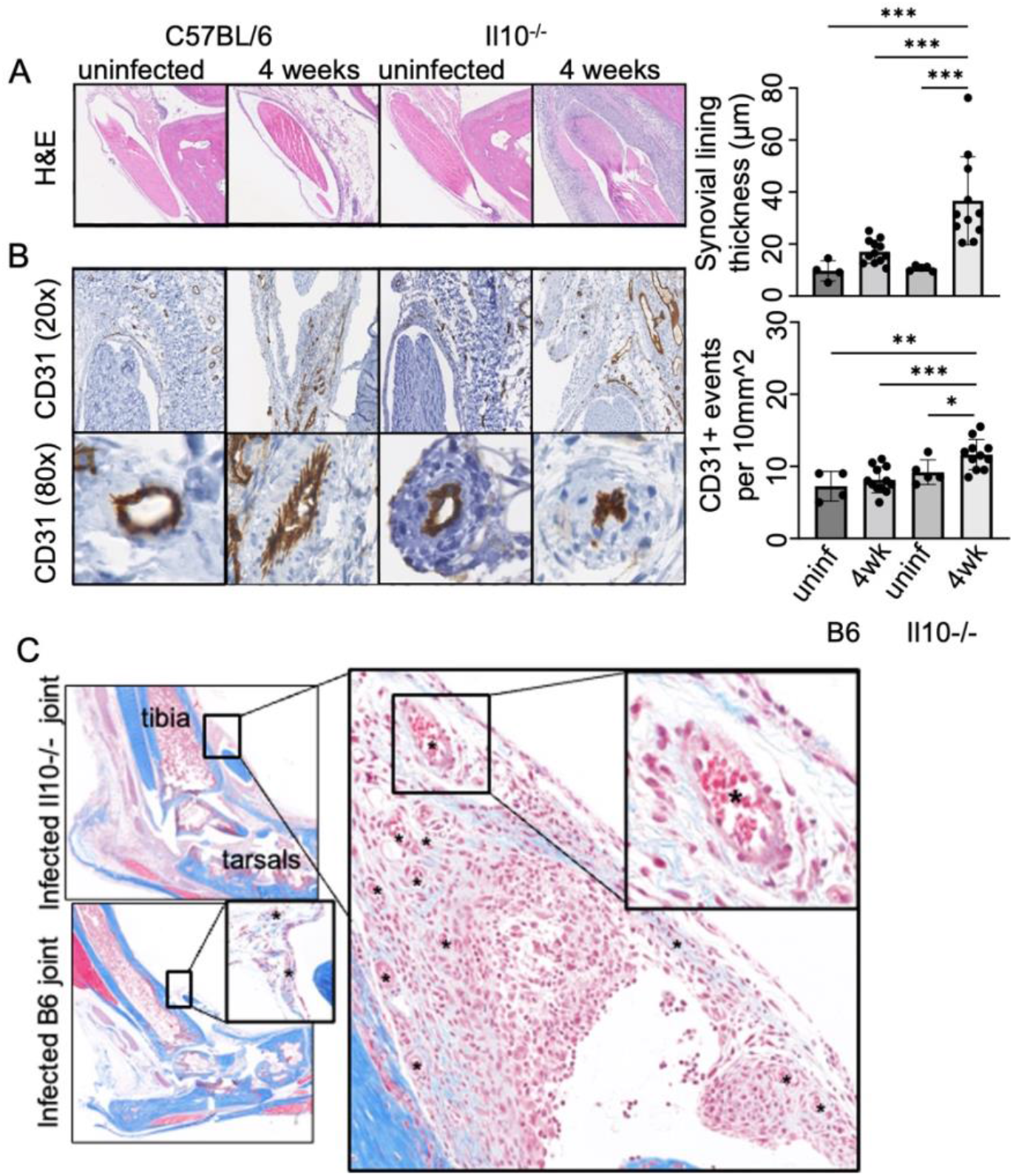
Synovial inflammation, fibrosis, and neovascularization in tibiotarsal joints of infected Il10^-/-^ mice. (A) The synovial lining surrounding the tibiotarsal tendon was measured (average of 5 measurements) from joint tissue stained with H&E. (B) Vascularization was determined by quantifying the number of CD31+ events per 10mm^2 using sections stained for PCAM (CD31) by immunohistochemistry. (C) Fibrosis was observed in synovial tissue in all infecdted mice by Masson’s trichrome staining. Collagen fibers (blue) can be seen within inflamed synovial tissue surrounding the tibiotarsal tendon. Blood vessels are labeled with an asterisk. No statistically significant differences in fibrosis were seen between groups. Representative images used for analysis are shown. Results are from 10 infected and 4-5 uninfected mice per group, statistically significance differences between groups were determined by Kruskal-Wallis discreet-variable analysis (*P<0.05, **P<0.01, ***P<0001).

### Identification of an immunogenic peptide derived from Borrelia *burgdorferi* Mcp4

Our immunopeptidomics analysis also identified several candidate peptides derived from *B. burgdorferi* proteins. Of these, six had spectra and retention times (RT) that matched the synthesized peptide (Mimitopes, Victoria, Australia). To determine whether any of these epitopes were immunogenic in mice, we infected B6 and Il10^-/-^ mice with *B. burgdorferi* for 4 weeks and tested for CD4+ T cell reactivity by stimulating lymphocytes with each peptide or a negative control peptide (**Fig 5A**). One of the six peptides elicited robust CD4+ T cell proliferation and activation, as measured by upregulation of CD69 in two of four Il10^-/-^ mice, but not in B6 mice (**Fig 5B**). This peptide was derived from BB0680, methyl-accepting chemotaxis protein 4 (Mcp4_442-462_), within the cytosolic methyl-accepting transducer domain (**Fig 5C**). To our knowledge, this is the first *B. burgdorferi-*derived MHC class II epitope identified in infected mice using this unbiased LC-MS/MS approach. Furthermore, these data indicate that IL-10 may play an important role in suppressing CD4+ T cell responses against *B. burgdoferi*, which has been previously suggested(28).

**Figure 5:**
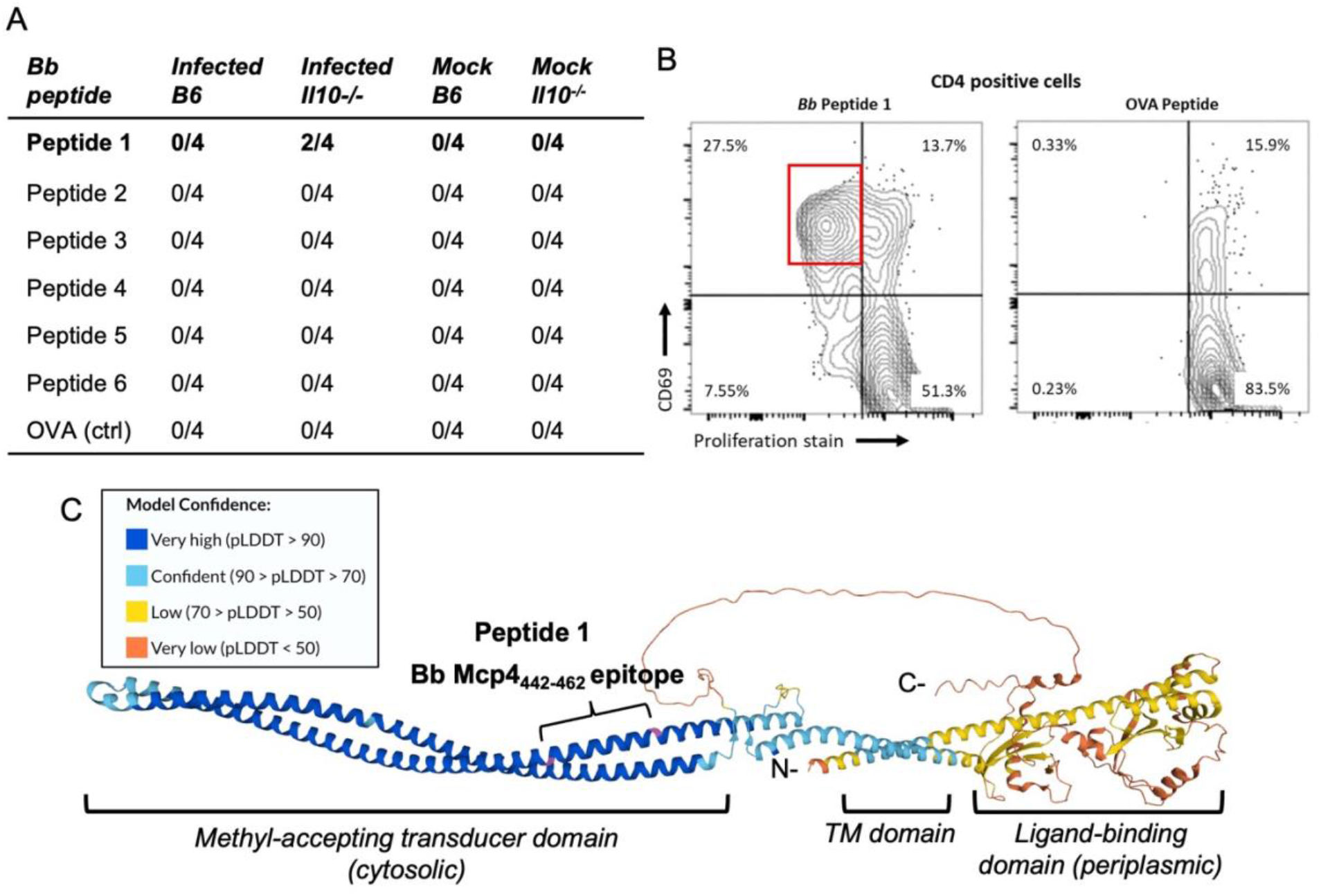
CD4+ T cell epitope Mcp4_442-462_ (Peptide 1) is immunogenic in Il10^-/-^ mice infected with *Bb*. (A) Lymphocytes from mice (4 animals per group) infected with *Bb* or BSKII (mock) for 4 weeks were stained with cell-trace violet (proliferation stain) and stimulated with *Bb* peptides 1-6 that were identified in our immunopeptidomics screen, or with OVA control peptide, for 5 days. (B) Flow cytometric identification of CD69+ (activated) proliferating population of CD4+ lymphocytes, highlighted with a red box, from a *Bb*-infected Il10^-/-^ mouse stimulated with Peptide 1 (Mcp4_442-462_) or OVA peptide control, gated on live singlet CD4+ lymphocytes. (C) Alpha-fold model of Mcp4 (BB0680) is shown with the localization of Peptide 1 in the methyl- accepting transducer domain.

## DISCUSSION

In this study, we used an immunopeptidomics approach to examine how the MHC class II peptide repertoire changes during *B. burgdorferi* infection in an effort to address critical gaps in our understanding of CD4+ T cell responses to *B. burgdorferi* antigens and to potential self-antigens in LD. Using methods developed to identify Lyme autoantigens in humans(64), we identified nearly 10,000 predicted peptides bound to MHC class II molecules isolated from inguinal and popliteal lymph nodes collected from mice infected with *B. burgdorferi* for 0, 4, or 16 weeks. Many of these were derived from proteins associated with extracellular matrix remodeling and vascular damage. Most notably apoB-100, a validated target of autoimmune T cell responses in human LA(16), cardiovascular disease(37), and liver damage(50), was encountered frequently. Infected mice showed evidence of marked apoB-100 epitope spreading and release of immunogenic cryptic autoantigens, which was most dramatic in mice lacking the immunosuppressive cytokine IL-10. Predicted peptides from *B. burgdorferi* proteins were synthesized and validated by LC-MS/MS, and six were identified as *bone fide* MHC class II epitopes. One *B. burgdorferi* epitope, derived from the predicted methyl-accepting chemotaxis protein MCP4 (BB0680), was found to be a CD4+ T cell immunogen in *B. burgdorferi*-infected mice lacking IL-10, but not in wild-type C57BL/6 (B6) mice. Thus, IL-10 appears to play a key role in regulating autoantigen epitope spreading and suppressing pathogen-specific T cell responses in mice infected with *B. burgdorferi*.

A key finding of this study was evidence of epitope spreading of apoB-100, a known human Lyme autoantigen, in *B. burgdorferi*-infected mice. Importantly, apoB-100 epitope spreading appeared to be tightly regulated by IL-10. Autoimmunity to apoB-100 in atherosclerosis can either be protective or pathogenic(37), and it is still unknown whether hyperlipidemia and apoB- 100 autoimmunity contribute to LD pathogenesis, particularly in the post-antibiotics stage.

Recent publications indicate, however, that high levels of cholesterol may exacerbate LD symptoms during *B. burgdorferi* infection(33) and contribute to PTLDS(32). Interestingly, hypercholesteremic ApoE^-/-^ and Ldlr^-/-^ mice on Western diets are prone to more severe LA and higher numbers of spirochetes in tissues, compared with their wild-type counterparts on normal chow(68). It is possible that apoB-100 reactive T cells may be limited in number or have an anti-inflammatory effect in most cases, but under certain conditions (e.g., individuals with elevated LDL levels), apoB-100 reactive T cells may become pro-inflammatory and pathogenic. Plasticity of apoB-100 T effector cells has has been demonstrated in human and murine atherosclerosis(37), supporting this hypothesis.

We hypothesize that epitope spreading originates from the point of attachment to LDLR from the self-tolerized P1 peptide towards either P2 or the other identified peptides within close proximity, some of which may become immunogenic in patients with LD. This region may be inherently immunogenic, particularly in mice and humans with elevated LDL levels(38). According to this hypothesis, we also predict that LD patients with hypercholesterolemia and/or hyperlipidemia will also have increased numbers of autoreactive apoB-100 T cells. Future studies will be required to test this hypothesis.

It is tempting to speculate that there is a causal link between *B. burgdorferi* acquisition of cholesterol and cholesterol-glycolipids during infection and apoB-100 epitiope spreading. *B. burgdorferi* inner and outer membranes consist of ∼45% cholesterol and cholesterol- glycolipids(59) and LDL particles are the most abundant sources of the precursor lipid molecules scavanged by *B. burgdorferi* for membrane synthesis. Membrane exchange has been demonstrated to occur between *B. burgdorferi* and mammalian cells(56), and apoB-100/LDL particles play a central role recycling of lipids from damaged cells and tissues that accumulate during vascular inflammation(36). Thus, the trigger for apoB-100 epitope spreading may be immune responses to *B. burgdorferi* glycolipids that are associated with LDL particles within the inflamed vasculature. NKT cells targeting *B. burgdorferi* glycolipids are highly activated during infection(62, 69). These activated NKT cells may provide the apoB-100 autoimmune trigger by promoting APCs to present *B. burgdorferi* lipid antigens to NKT cells and apoB-100 epitopes to potentially self-reactive T cells, likely involving Kuppfer cells and NKT cells within the liver(70).

The second key finding of this study was identification of Mcp4_442-462_ as a *bone fide* T cell immunogen in mice. There is a paucity of known CD4+ T cell epitopes derived from *B. burgdorferi* proteins, which has significantly hampered studies of T cell immune responses during *B. burgdorferi* infection(25). T cell responses to Mcp4 appear to be suppressed by IL-10. This may play an important role in *B. burgdorferi* evasion of adaptive immune responses, which can now be explored using Mcp4_442-462_ as a model antigen.

It is intriguing that apoB-100 epitope spreading and Mcp4 immunoreactivity is regulated by IL-10. This cytokine has been well-studied in murine LD and is believed to be a key regulator of both inflammation and host defense in mice and humans(4, 28, 30, 52, 71). This study supports this hypothesis, particularly as it relates to CD4+ T cell responses to foreign and self-antigens during infection, and provides novel immune targets that can be exploited to elucidate mechanisms of immune evasion and development of infection-induced autoimmunity in patients with Lyme disease. Future studies will be needed to test this hypothesis.

An inherent limitation of this mouse model study is that LA and T cell autoimmunity to apoB-100 takes ∼6-12 months to develop in humans(11, 16). Therefore, it is challenging to study the entire process of infection-induced autoimmunity in mice, which develop LA at ∼4 weeks post- infection, and mice and humans have markedly different compositions of lipoproteins, including LDL(72). Despite this limitation, use of the mouse models in this study allowed us to examine immune processes that occur early during infection in an experimentally tractable system to gain insight into the initial steps in a path that eventually leads to autoimmunity in humans. It is likely that additional studies using different mouse models, such as the hypercholesterolemic ApoE^-/-^ or Ldlr^-/-^ mouse models, will be needed to gain further insights into the effects of B. burgdorferi infection on apoB-100 autoimmunity.

## MATERIALS AND METHODS

### Ethics statement

Mice were housed in the Medical College of Wisconsin Biomedical Resource Center (Milwaukee, WI), following strict adherence to the guidelines according to the National Institutes of Health for the care and use of laboratory animals, as described In the Guide for the Care and Use of Laboratory Animals, 8^th^ Edition. Protocols conducted in this study were approved and carried out in accordance with the Medical College of Wisconsin Institutional Animal Care and Use Committee (IACUC protocol number AUA00006528). Mouse experiments were performed under isoflurane anesthesia, and every effort was made to minimize suffering. Research design and outcome measures were selected according to ARRIVE 2.0 guidelines(73) to ensure that reviewers and readers can assess the reliability of the findings presented.

### Mouse infection and lymph node preparation

C57BL/6 and B6.129P2-IL-10^tm1Cgn^/J (B6 Il10^−/−^) mice were obtained from the Jackson Laboratory and injected with 0.02 mL of either 2×10^3^ *B. burgdorferi* (strain N40) or BSKII medium (negative control) by intradermal injection into the skin of the back. At 4- and 16-weeks post-inoculation, inguinal and popliteal lymph nodes were harvested and pooled per mouse group and stored in liquid nitrogen. On the day of purification, 0.6 – 1.8 g of tissue was thawed and processed using a BeadBug-6 Homogenizer (Benchmark Scientific) on ice with 12 mL of lysis buffer (150 mM sodium chloride, 20 mM tris-(hydroxymethyl)aminomethane·HCl (pH 8.0), 5 mM EDTA solution, 0.04% sodium azide, 1 mM 4-(2-aminoethyl)benzenesulfonyl fluoride·HCl, 10 μg/mL leupeptin, 10 μg/mL pepstatin A, 5 μg/mL aprotinin, and 1% 3-[(3- cholamidopropyl)dimethylammonio]-1-propanesulfonate). The detergent CHAPS was then added to 1% final concentration, and the extract was incubated at 4°C for 1 h. Insoluble material was removed by centrifugation at 800 × g for 5 min and 27,000 × g for 30 min.

### Immunoaffinity purification of MHC-II-peptide complexes

The procedures used for MHC II peptide isolation were similar to those published by Seward, *et al* (31). Water (Fisher Scientific) was HPLC grade. CNBr-activated Sepharose beads (GE Healthcare) were used for coupling of the I-A/I-E-specific antibody (M5/114.15.2; 4 mg). Before antibody coupling, the Sepharose beads were prehydrolyzed in coupling buffer (0.1M NaHCO3, 0.5M NaCl, pH 8.5) to reduce the number of reactive groups on the beads. After prehydrolysis, antibody was incubated with the beads for 1 h followed by a 3 h incubation with blocking buffer (0.015 g/mL glycine in coupling buffer). Uncoupled beads were prepared identically but without antibody. Cell lysates were incubated with uncoupled beads for 4 h at 4°C and then with M5/114.15.2-conjugated beads overnight at 4°C. Beads were washed 4 × 15 min and 1 × 30 min with 14 mL of 1% CHAPS-containing lysis buffer and then 4 × 15 min and 1 × 40 min with 14 mL of High NaCl wash buffer (150 mM NaCl, 20 mM Tris-HCl, pH 8). Beads were transferred to a column (5 mL, Pierce) and washed with 50 mL of High NaCl wash buffer followed by 50 mL of No NaCl wash buffer (20 mM Tris-HCl, pH 8) by gravity flow. Residual buffer was removed by centrifugation for 2 min at 800 × g. Peptides were eluted by 7 × 5 min incubations at RT with 0.7 mL of 0.5% formic acid followed by centrifugation for 2 min at 800 × g to collect eluates.

### LC-MS/MS

Peptide extracts were dried under vaccum and re-dissolved in 28 µL of 2% acetonitrile/0.1% formic acid with vortexing and sonication. After centrifugation, 24 µL of the samples were transferred to autosampler vials for duplicate injections of 10 µl onto a 25 cm C18 column (Microm Bioresources Magic C18AQ). The peptides were then separated using a 90-minute method and the subsequently analyzed on a Thermo Orbitrap Fusion Lumo MS via two technical replicate ingections using a data-dependent acquisition (DDA) HCD MS2 instrument method.

### Protein Database Searching

MS data were analyzed using the Proteome Discoverer 2.4 (Thermo) platform. SequestHT was used as search algorithm. Percolator and Protein FDR Validator were used for validation. Proteome databases searched for this study were *Borrelia burgdorferi* (N40 and B31), mouse (Swissprot mouse, with isoforms), and MaxQuant contaminants. Oxidation (M), acetylation (protein N-terminus), deamidation (N, Q), and Gln->pyro-Glu (peptide N-terminus) dynamic modifications were also assessed. Target FDR for PSMs and peptides were 0.01 (strict) or 0.05 (relaxed).

### Validation of *B. burgdorferi* peptides

Candidate peptides identified by LC-MS/MS were synthesized by Mimitopes (Victoria, Australia). Purified peptides were analyzed by LC-MS/MS as above, and mass spectra and retention times (RT) were compared. Six *B. burgdorferi* peptides with matching spectra and RT were selected for further testing.

### Histology

Histology sectioning and imaging were performed by the Children’s Hospital of Wisconsin Histology Core Facility. Tibiotarsal joints were harvested from B6 and Il10^-/-^ mice and stored in 10% formalin (Sigma Aldrich) for 72 hours. Samples were decalcified, embedded into paraffin, sectioned, and stained with hematoxylin and eosin (H&E), Masson’s trichrome, and anti-CD31. Two sections were analyzed for each joint, and special care was made to ensure that each section was obtained from the center portion of the joint.

Two joints from each mouse were used to assess inflammation by H&E staining and neovascularization by anti-CD31 staining at 10X magnification. The synovial lining thickness surrounding the tibiotarsal tendon was recorded from five, random measurements for each joint. The number of CD31-stained events were recorded from five, random 10mm^2^ areas of the synovium surrounding the tibiotarsal tendon. The average of two joints per mouse were plotted by group for analysis of inflammation and neovascularization.

### T-cell proliferation assay

APCs and T cells were isolated from inguinal lymph nodes of 4-week infected C57BL/6 mice and B6 IL10^−/−^ mice. Cells were cultured in complete RPMI medium (10% FBS, 2 mM L-glutamine, 100 U/mL penicillin, 100 µg/mL streptomycin, 1% nonessential amino acids) with IL-2 (0.3125 µg/mL) at 1x10^5^ cells/well in a 96-well plate. Cells were stained with 5 µM of CellTrace Violet (Invitrogen) according to the manufacturer’s instructions, stimulated with 2 µM of each peptide (Mimitopes) or 2.4 µg/mL of OVA_257-264_ peptide (negative control), and incubated at 37°C/5% CO_2_ for 5 days. Cells were collected and washed with phosphate-buffered saline (PBS) before incubation with cell surface antibodies (BioLegend; CD4-PE, CD69-APC, and CD90.1-PE/Cy7) for 20 minutes. Cells were analyzed using FACS Celesta (Beckman Coulter) and FlowJo v.10.7.1 software.

### Protein structure modeling

The *in silico*-generated protein structure of human apoB-100 was generously provided by Jeiran, *et al* (74). Their group separated apoB-100 into five separate domains, used multiple protein modeling programs to construct energy minimized and refined models of each domain, and finally processed the individual models within AlphaFold before joining them together into the complete ∼4.6 kDa protein. As an added measure of validation, the group synthesized portions of the protein in conjunction with the mass spectrometry-cleavable cross-linker disuccinimidyl sulfoxide (DSSO) to determine the position of disulfide bonds within apoB-100 when joining the separate domains. Approximately 88% of the 65 unique DSSO cross-links were within the expected 26 Å that these tertiary bonds are known to span, as determined by nanoscale LC-MS/MS. The subsequent file containing the complete apoB-100 structure was then analyzed using PyMol v2.5.4 (The PyMOL Molecular Graphics System, Version 2.0 Schrödinger, LLC.) to locate and highlight the MHC-II presented peptides as identified from the LC-MS/MS data. Presented peptides were derrived from mice, however the amino acid sequence homology between human and murine apoB-100 is approximately 70% identical. To account for any species-specific alterations in amino acid sequence, the peptides were searched for within the sequence allowing for up to five mismatches.

## ACKNOWLEDGEMENTS

We would like to thank Kianoush Jeiran, MD, PhD, (NHLBI and George Mason University) and Alan Remaley, MD, PhD, (NHLBI) for generously providing the apoB-100 pdb file used to generate the structural model. We would also like to acknowledge Christine Duris, BS HTL QIHC (ASCP), and the other members of the MCW Childrens Research Institute Histology Core for their invaluable assistance in preparing and staining tissue sections.

## SUPPLEMENTAL MATERIAL

**Table 1.**
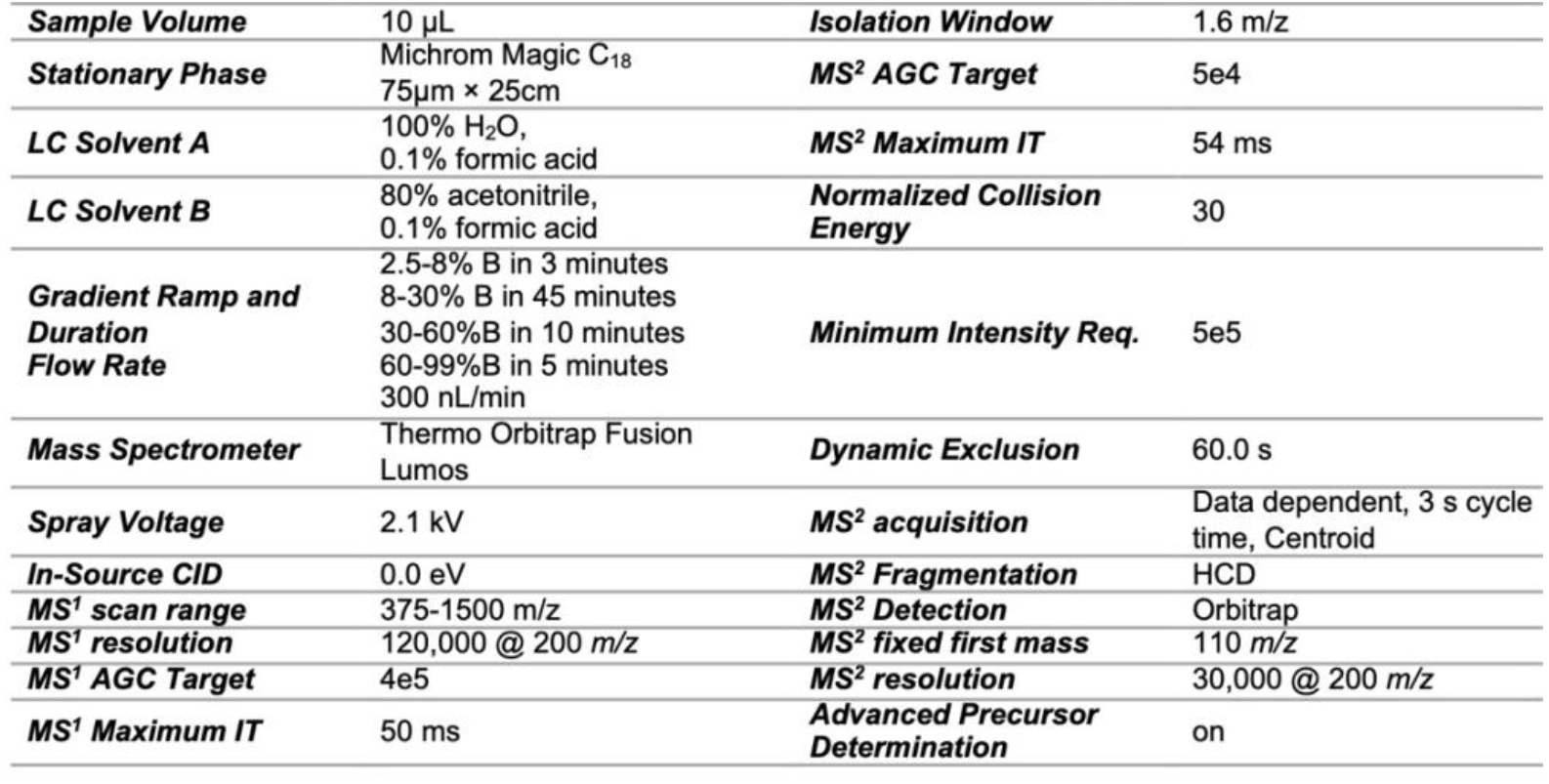
Chromatography and MS instrument acquisition settings.

**Table 2.** Mass spectrometry data processing parameters.

| Table 2. Mass spectrometry data processing parameters |  |  |  |
| --- | --- | --- | --- |
| **Platform** | ProteomeDiscoverer 2.4 | **Quantitation** |  |
| **Search Algorithms** | SequestHT | **Normalization** |  |
| **Validation** | Percolator, Protein FDR Validator | **Scaling** |  |
| **Database** | Borrelia burgdorferi B31<br>2019-09-25 | **Ratio Calculation** |  |
|  | Borrelia burgdorferi N40<br>2019-09-25 |  |  |
|  | Swissprot mouse with isoforms, 2019-05-01,<br>MaxQuant Contaminants |  |  |
| **Digest** | No-Enzyme (Unspecific) | **Hypothesis Test** |  |
| **Precursor mass tolerance** | 10 ppm | **Target FDR (Strict) for PSMs:** | 0.01 |
| **Fragment mass tolerance** | 0.02 Da | **Target FDR (Relaxed) for PSMs:** | 0.05 |
| **Static Modifications** |  | **Target FDR (Strict) for Peptides:** | 0.01 |
| **Dynamic Modifications** | Oxidation (M), Acetylation (protein N-terminus)<br>Deamidation (N,Q)<br>Gln→pyro-Glu | **Target FDR (Relaxed) for Peptides:** | 0.05 |

**Supplemental Figure S1.**
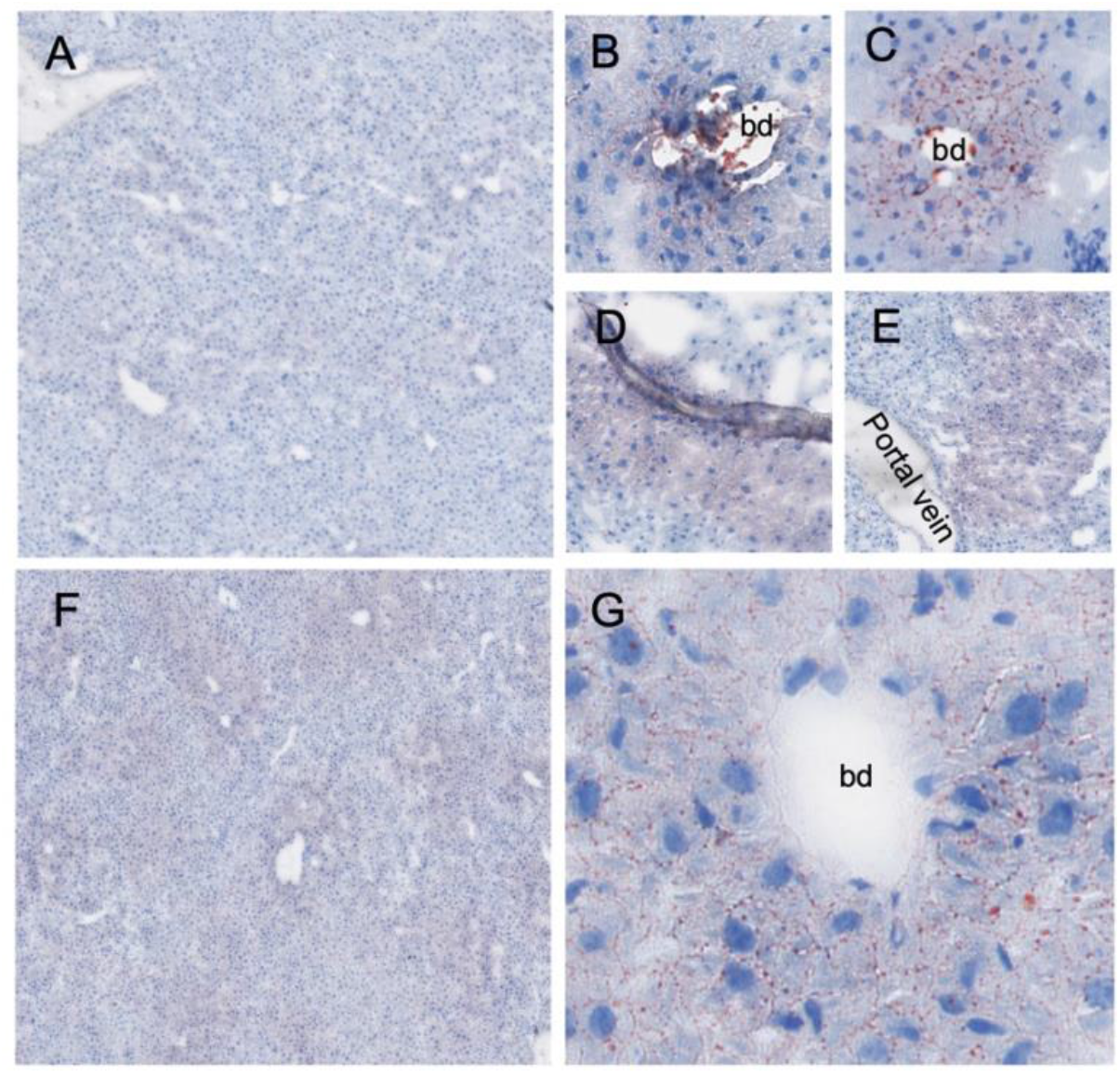
Neutral lipid deposits in livers from B6 and 1110-/- mice infected with *Bb.* (A) Liver sections from a mock-infected B6 and 1110-/- mice fed normal chow has minimal accumulation of triglycerides and other neutral lipids, as measured by Oil Red O staining. B6 and 1110-/- mice infected with *Bb* for 1 week typically had 0-4 regions within each liver section that stain positive for neutral lipids (red, B-E). These regions often surround bile ducts (bd, B-C), which may either have modest inflammation (B) or little to no inflammation (C); or the small (D) or large (E) vessels of the liver. (F) One infected 1110-/- mouse had mild, diffuse accumulation of neutral lipids, which was most abundant surrounding bile ducts (enlarged in G). The other infected 1110-/- mice had small, localized areas of lipid accumulation, like those shown in B-E. There were no statistically significant differences in lipid deposition between strains. Images are representative of 10 βfc>-infected mice and 4 uninfected mice per strain, 3 sections per mouse.

## REFERENCES

1. Cooper GS, Bynum ML, Somers EC. Recent insights in the epidemiology of autoimmune diseases: improved prevalence estimates and understanding of clustering of diseases. J Autoimmun. 2009;33(3-4):197–207.

2. Bach JF. The effect of infections on susceptibility to autoimmune and allergic diseases. N Engl J Med. 2002;347(12):911–20.

3. von Herrath MG, Fujinami RS, Whitton JL. Microorganisms and autoimmunity: making the barren field fertile? Nat Rev Microbiol. 2003;1(2):151–7.

4. Lochhead RB, Strle K, Arvikar SL, Weis JJ, Steere AC. Lyme arthritis: linking infection, inflammation and autoimmunity. Nat Rev Rheumatol. 2021;17(8):449–61.

5. Dong Y, Zhou G, Cao W, Xu X, Zhang Y, Ji Z, et al. Global seroprevalence and sociodemographic characteristics of Borrelia burgdorferi sensu lato in human populations: a systematic review and meta-analysis. BMJ Glob Health. 2022;7(6).

6. Kugeler KJ, Schwartz AM, Delorey MJ, Mead PS, Hinckley AF. Estimating the Frequency of Lyme Disease Diagnoses, United States, 2010-2018. Emerg Infect Dis. 2021;27(2):616–9.

7. Radolf JD, Caimano MJ, Stevenson B, Hu LT. Of ticks, mice and men: understanding the dual-host lifestyle of Lyme disease spirochaetes. Nat Rev Microbiol. 2012;10(2):87–99.

8. Steere AC, Strle F, Wormser GP, Hu LT, Branda JA, Hovius JW, et al. Lyme borreliosis. Nat Rev Dis Primers. 2016;2:16090.

9. Mead PS. Epidemiology of Lyme disease. Infect Dis Clin North Am. 2015;29(2):187–210.

10. Sanchez JL. Clinical Manifestations and Treatment of Lyme Disease. Clin Lab Med. 2015;35(4):765–78.

11. Arvikar SL, Steere AC. Lyme Arthritis. Infect Dis Clin North Am. 2022;36(3):563–77.

12. Bobe JR, Jutras BL, Horn EJ, Embers ME, Bailey A, Moritz RL, et al. Recent Progress in Lyme Disease and Remaining Challenges. Front Med (Lausanne). 2021;8:666554.

13. Aucott JN. Posttreatment Lyme disease syndrome. Infect Dis Clin North Am. 2015;29(2):309–23.

14. Steere AC. Posttreatment Lyme disease syndromes: distinct pathogenesis caused by maladaptive host responses. J Clin Invest. 2020;130(5):2148–51.

15. Marques A. Persistent Symptoms After Treatment of Lyme Disease. Infect Dis Clin North Am. 2022;36(3):621–38.

16. Crowley JT, Drouin EE, Pianta A, Strle K, Wang Q, Costello CE, et al. A Highly Expressed Human Protein, Apolipoprotein B-100, Serves as an Autoantigen in a Subgroup of Patients With Lyme Disease. J Infect Dis. 2015;212(11):1841–50.

17. Crowley JT, Strle K, Drouin EE, Pianta A, Arvikar SL, Wang Q, et al. Matrix metalloproteinase-10 is a target of T and B cell responses that correlate with synovial pathology in patients with antibiotic-refractory Lyme arthritis. J Autoimmun. 2016;69:24–37.

18. Pianta A, Drouin EE, Crowley JT, Arvikar S, Strle K, Costello CE, et al. Annexin A2 is a target of autoimmune T and B cell responses associated with synovial fibroblast proliferation in patients with antibiotic-refractory Lyme arthritis. Clin Immunol. 2015;160(2):336–41.

19. Drouin EE, Seward RJ, Strle K, McHugh G, Katchar K, Londono D, et al. A novel human autoantigen, endothelial cell growth factor, is a target of T and B cell responses in patients with Lyme disease. Arthritis Rheum. 2013;65(1):186–96.

20. Brouwer MAE, van de Schoor FR, Vrijmoeth HD, Netea MG, Joosten LAB. A joint effort: The interplay between the innate and the adaptive immune system in Lyme arthritis. Immunol Rev. 2020;294(1):63–79.

21. Elsner RA, Hastey CJ, Olsen KJ, Baumgarth N. Suppression of Long-Lived Humoral Immunity Following Borrelia burgdorferi Infection. PLoS Pathog. 2015;11(7):e1004976.

22. Tracy KE, Baumgarth N. Borrelia burgdorferi Manipulates Innate and Adaptive Immunity to Establish Persistence in Rodent Reservoir Hosts. Front Immunol. 2017;8:116.

23. Hammond EM, Baumgarth N. CD4 T cell responses in persistent Borrelia burgdorferi infection. Curr Opin Immunol. 2022;77:102187.

24. Brouwer MAE, Jones-Warner W, Rahman S, Kerstholt M, Ferreira AV, Oosting M, et al. B. burgdorferi sensu lato-induced inhibition of antigen presentation is mediated by RIP1 signaling resulting in impaired functional T cell responses towards Candida albicans. Ticks Tick Borne Dis. 2021;12(2):101611.

25. Helble JD, McCarthy JE, Sawden M, Starnbach MN, Hu LT. The PD-1/PD-L1 pathway is induced during Borrelia burgdorferi infection and inhibits T cell joint infiltration without compromising bacterial clearance. PLoS Pathog. 2022;18(10):e1010903.

26. Elsner RA, Hastey CJ, Baumgarth N. CD4+ T cells promote antibody production but not sustained affinity maturation during Borrelia burgdorferi infection. Infect Immun. 2015;83(1):48–56.

27. Barthold SW, Beck DS, Hansen GM, Terwilliger GA, Moody KD. Lyme borreliosis in selected strains and ages of laboratory mice. J Infect Dis. 1990;162(1):133–8.

28. Sonderegger FL, Ma Y, Maylor-Hagan H, Brewster J, Huang X, Spangrude GJ, et al. Localized production of IL-10 suppresses early inflammatory cell infiltration and subsequent development of IFN-gamma-mediated Lyme arthritis. J Immunol. 2012;188(3):1381–93.

29. Lochhead RB, Sonderegger FL, Ma Y, Brewster JE, Cornwall D, Maylor-Hagen H, et al. Endothelial cells and fibroblasts amplify the arthritogenic type I IFN response in murine Lyme disease and are major sources of chemokines in Borrelia burgdorferi-infected joint tissue. J Immunol. 2012;189(5):2488–501.

30. Whiteside SK, Snook JP, Ma Y, Sonderegger FL, Fisher C, Petersen C, et al. IL-10 Deficiency Reveals a Role for TLR2-Dependent Bystander Activation of T Cells in Lyme Arthritis. J Immunol. 2018;200(4):1457–70.

31. Seward RJ, Drouin EE, Steere AC, Costello CE. Peptides presented by HLA-DR molecules in synovia of patients with rheumatoid arthritis or antibiotic-refractory Lyme arthritis. Mol Cell Proteomics. 2011;10(3):M110 002477.

32. Chung MK, Caboni M, Strandwitz P, D’Onofrio A, Lewis K, Patel CJ. Systematic comparisons between Lyme disease and post-treatment Lyme disease syndrome in the U.S. with administrative claims data. EBioMedicine. 2023;90:104524.

33. Forrest IS, O’Neal AJ, Pedra JHF, Do R. Cholesterol contributes to risk, severity, and machine learning-driven diagnosis of Lyme disease. Clin Infect Dis. 2023.

34. Horowitz HW, Dworkin B, Forseter G, Nadelman RB, Connolly C, Luciano BB, et al. Liver function in early Lyme disease. Hepatology. 1996;23(6):1412–7.

35. Fitzgerald BL, Molins CR, Islam MN, Graham B, Hove PR, Wormser GP, et al. Host Metabolic Response in Early Lyme Disease. J Proteome Res. 2020;19(2):610–23.

36. Mehta A, Shapiro MD. Apolipoproteins in vascular biology and atherosclerotic disease. Nat Rev Cardiol. 2022;19(3):168–79.

37. Wolf D, Gerhardt T, Winkels H, Michel NA, Pramod AB, Ghosheh Y, et al. Pathogenic Autoimmunity in Atherosclerosis Evolves From Initially Protective Apolipoprotein B(100)- Reactive CD4(+) T-Regulatory Cells. Circulation. 2020;142(13):1279–93.

38. Fredrikson GN, Lindholm MW, Ljungcrantz I, Soderberg I, Shah PK, Nilsson J. Autoimmune responses against the apo B-100 LDL receptor-binding site protect against arterial accumulation of lipids in LDL receptor deficient mice. Autoimmunity. 2007;40(2):122–30.

39. Herbin O, Ait-Oufella H, Yu W, Fredrikson GN, Aubier B, Perez N, et al. Regulatory T- cell response to apolipoprotein B100-derived peptides reduces the development and progression of atherosclerosis in mice. Arterioscler Thromb Vasc Biol. 2012;32(3):605–12.

40. Nilsson J, Bjorkbacka H, Fredrikson GN. Apolipoprotein B100 autoimmunity and atherosclerosis - disease mechanisms and therapeutic potential. Curr Opin Lipidol. 2012;23(5):422–8.

41. Hermansson A, Ketelhuth DF, Strodthoff D, Wurm M, Hansson EM, Nicoletti A, et al. Inhibition of T cell response to native low-density lipoprotein reduces atherosclerosis. J Exp Med. 2010;207(5):1081–93.

42. Tse K, Gonen A, Sidney J, Ouyang H, Witztum JL, Sette A, et al. Atheroprotective Vaccination with MHC-II Restricted Peptides from ApoB-100. Front Immunol. 2013;4:493.

43. Shaw MK, Tse KY, Zhao X, Welch K, Eitzman DT, Thipparthi RR, et al. T-Cells Specific for a Self-Peptide of ApoB-100 Exacerbate Aortic Atheroma in Murine Atherosclerosis. Front Immunol. 2017;8:95.

44. Ley K. 2015 Russell Ross Memorial Lecture in Vascular Biology: Protective Autoimmunity in Atherosclerosis. Arterioscler Thromb Vasc Biol. 2016;36(3):429–38.

45. Kimura T, Kobiyama K, Winkels H, Tse K, Miller J, Vassallo M, et al. Regulatory CD4(+) T Cells Recognize Major Histocompatibility Complex Class II Molecule-Restricted Peptide Epitopes of Apolipoprotein B. Circulation. 2018;138(11):1130–43.

46. Kimura T, Tse K, McArdle S, Gerhardt T, Miller J, Mikulski Z, et al. Atheroprotective vaccination with MHC-II-restricted ApoB peptides induces peritoneal IL-10-producing CD4 T cells. Am J Physiol Heart Circ Physiol. 2017;312(4):H781–H90.

47. Klingenberg R, Lebens M, Hermansson A, Fredrikson GN, Strodthoff D, Rudling M, et al. Intranasal immunization with an apolipoprotein B-100 fusion protein induces antigen-specific regulatory T cells and reduces atherosclerosis. Arterioscler Thromb Vasc Biol. 2010;30(5):946–52.

48. Wolf D, Ley K. Immunity and Inflammation in Atherosclerosis. Circ Res. 2019;124(2):315–27.

49. Chyu KY, Zhao X, Zhou J, Dimayuga PC, Lio NW, Cercek B, et al. Immunization using ApoB-100 peptide-linked nanoparticles reduces atherosclerosis. JCI Insight. 2022;7(11).

50. Plochg BFJ, Englert H, Rangaswamy C, Konrath S, Malle M, Lampalzer S, et al. Liver damage promotes pro-inflammatory T-cell responses against apolipoprotein B-100. J Intern Med. 2022;291(5):648–64.

51. Duray PH, Steere AC. Clinical pathologic correlations of Lyme disease by stage. Ann N Y Acad Sci. 1988;539:65–79.

52. Lochhead RB, Zachary JF, Dalla Rosa L, Ma Y, Weis JH, O’Connell RM, et al. Antagonistic Interplay between MicroRNA-155 and IL-10 during Lyme Carditis and Arthritis. PLoS One. 2015;10(8):e0135142.

53. Lochhead RB, Ma Y, Zachary JF, Baltimore D, Zhao JL, Weis JH, et al. MicroRNA-146a provides feedback regulation of lyme arthritis but not carditis during infection with Borrelia burgdorferi. PLoS Pathog. 2014;10(6):e1004212.

54. Ordonez D, Lochhead RB, Strle K, Pianta A, Arvikar S, Van Rhijn I, et al. Cell-Mediated Cytotoxicity in Lyme Arthritis. Arthritis Rheumatol. 2023;75(5):782–93.

55. Casselli T, Divan A, Vomhof-DeKrey EE, Tourand Y, Pecoraro HL, Brissette CA. A murine model of Lyme disease demonstrates that Borrelia burgdorferi colonizes the dura mater and induces inflammation in the central nervous system. PLoS Pathog. 2021;17(2):e1009256.

56. Crowley JT, Toledo AM, LaRocca TJ, Coleman JL, London E, Benach JL. Lipid exchange between Borrelia burgdorferi and host cells. PLoS Pathog. 2013;9(1):e1003109.

57. O’Neal AJ, Butler LR, Rolandelli A, Gilk SD, Pedra JH. Lipid hijacking: a unifying theme in vector-borne diseases. Elife. 2020;9.

58. Gwynne PJ, Clendenen LH, Turk SP, Marques AR, Hu LT. Antiphospholipid autoantibodies in Lyme disease arise after scavenging of host phospholipids by Borrelia burgdorferi. J Clin Invest. 2022;132(6).

59. Radolf JD, Goldberg MS, Bourell K, Baker SI, Jones JD, Norgard MV. Characterization of outer membranes isolated from Borrelia burgdorferi, the Lyme disease spirochete. Infect Immun. 1995;63(6):2154–63.

60. Jones KL, Seward RJ, Ben-Menachem G, Glickstein LJ, Costello CE, Steere AC. Strong IgG antibody responses to Borrelia burgdorferi glycolipids in patients with Lyme arthritis, a late manifestation of the infection. Clin Immunol. 2009;132(1):93–102.

61. Kumar H, Belperron A, Barthold SW, Bockenstedt LK. Cutting edge: CD1d deficiency impairs murine host defense against the spirochete, Borrelia burgdorferi. J Immunol. 2000;165(9):4797–801.

62. Szamosvari D, Bae M, Bang S, Tusi BK, Cassilly CD, Park SM, et al. Lyme Disease, Borrelia burgdorferi, and Lipid Immunogens. J Am Chem Soc. 2022;144(6):2474–8.

63. Vanderlugt CL, Miller SD. Epitope spreading in immune-mediated diseases: implications for immunotherapy. Nat Rev Immunol. 2002;2(2):85–95.

64. Wang Q, Drouin EE, Yao C, Zhang J, Huang Y, Leon DR, et al. Immunogenic HLA-DR- Presented Self-Peptides Identified Directly from Clinical Samples of Synovial Tissue, Synovial Fluid, or Peripheral Blood in Patients with Rheumatoid Arthritis or Lyme Arthritis. J Proteome Res. 2017;16(1):122–36.

65. Rosenblum MD, Remedios KA, Abbas AK. Mechanisms of human autoimmunity. J Clin Invest. 2015;125(6):2228–33.

66. Herijgers N, Van Eck M, Groot PH, Hoogerbrugge PM, Van Berkel TJ. Low density lipoprotein receptor of macrophages facilitates atherosclerotic lesion formation in C57Bl/6 mice. Arterioscler Thromb Vasc Biol. 2000;20(8):1961–7.

67. Covarrubias R, Wilhelm AJ, Major AS. Specific deletion of LDL receptor-related protein on macrophages has skewed in vivo effects on cytokine production by invariant natural killer T cells. PLoS One. 2014;9(7):e102236.

68. Toledo A, Monzon JD, Coleman JL, Garcia-Monco JC, Benach JL. Hypercholesterolemia and ApoE deficiency result in severe infection with Lyme disease and relapsing-fever Borrelia. Proc Natl Acad Sci U S A. 2015;112(17):5491–6.

69. Katchar K, Drouin EE, Steere AC. Natural killer cells and natural killer T cells in Lyme arthritis. Arthritis Res Ther. 2013;15(6):R183.

70. Lee WY, Moriarty TJ, Wong CH, Zhou H, Strieter RM, van Rooijen N, et al. An intravascular immune response to Borrelia burgdorferi involves Kupffer cells and iNKT cells. Nat Immunol. 2010;11(4):295–302.

71. Lochhead RB, Strle K, Kim ND, Kohler MJ, Arvikar SL, Aversa JM, et al. MicroRNA Expression Shows Inflammatory Dysregulation and Tumor-Like Proliferative Responses in Joints of Patients With Postinfectious Lyme Arthritis. Arthritis Rheumatol. 2017;69(5):1100–10.

72. Gordon SM, Li H, Zhu X, Shah AS, Lu LJ, Davidson WS. A comparison of the mouse and human lipoproteome: suitability of the mouse model for studies of human lipoproteins. J Proteome Res. 2015;14(6):2686–95.

73. Percie du Sert N, Ahluwalia A, Alam S, Avey MT, Baker M, Browne WJ, et al. Reporting animal research: Explanation and elaboration for the ARRIVE guidelines 2.0. PLoS Biol. 2020;18(7):e3000411.

74. Jeiran K, Gordon SM, Sviridov DO, Aponte AM, Haymond A, Piszczek G, et al. A New Structural Model of Apolipoprotein B100 Based on Computational Modeling and Cross Linking. Int J Mol Sci. 2022;23(19).

